# Electrical Impedance Spectroscopy to easily assess structural changes in dominant versus auxiliary upper arms due to pathophysiological events

**DOI:** 10.1101/622944

**Authors:** A. H. Dell’Osa, A. Fois, Q. Mela, A. Loviselli, A. Capone, G. Marongiu, Filippo Tocco, A. Concu, F. Velluzzi

**Affiliations:** Instituto de Desarrollo Económico e Innovación, Universidad Nacional de Tierra del Fuego, Ushuaia, Argentina; Biosingals Acquisition Unit, Nomadyca Ltd, Kampala, Uganda; Osteoporosis Center, Department of Medical Sciences and Public Health, University of Cagliari, Italy; Obesity Center, Department of Medical Sciences and Public Health, University of Cagliari, Italy; Orthopaedic Clinic, Department of Surgical Sciences, University of Cagliari, Italy; Sports Medicine, Department of Medical Sciences and Public Health, University of Cagliari, Italy; 2C Technologies Ltd, Academic Spin-Off, University of Cagliari, Italy

**Keywords:** Electrical impedance spectroscopy, forearms electrical resistance, arms lymphedema, AD 5933 electronic board, Bland and Altman plots

## Abstract

The resistive component of the bioimpedance was non invasively assessed in both right and left forearms of 11 healthy female and 9 males (28.4 ± 1.4 years; 63.8 ± 11.8 kg; 167.4 ± 7.5 cm) all of which were right-handed. A homemade electrical impedance spectroscopy device which implemented the AD 5933 electronic board from Analog Devices Inc., USA, was utilized, and the bipolar modality of bioimpedance assessment was chosen using two disposable ECG surface electrodes placed in each end of the biceps brachial muscles while subject were comfortably sitting. Forearms resistance was acquired at sweeping frequencies steps of 15, 30, 45, 60 and 75 KHz. Results showed a significantly lower man value of resistance in right versus left forearms (- 27.4 Ω, P<0.05), or about -4%, at the frequency of 15 KHz. Even though there was a progressive reduction, this right versus left forearm resistance difference persisted as statistically significant up to the frequency of 45 KHz. It was concluded that the risk of some mistakes do exists when lymphedema may occur in one arm and electrical impedance spectroscopy was utilized to monitoring in that arm the water volume trend in comparison with the other side arm since these results underline in the main forearm a largely low value of the resistance than in the auxiliary one, even in healthy subjects. So, care must be taken when the electrical impedance spectroscopy was adopted in these clinical assessments.

## I. Introduction

Changes in the structural composition of the arm of one side with respect to that of the other side often may indicate not only proper adaptations in that arm, like as what happens in athletes engaged in asymmetric sports specialties [1], but also several illness occurring in other body organs. In fact, it has been found that a lot of post –breast cancer people shows accumulation of lymph fluid in the arm of the affected side [2]. In considering that it is well known that a strict inverse correlation does exist between the water content in a body segment and the real component of the electrical impedance (i.e. the resistance) when an alternate electrical current flows through that body at a low frequency [3], it has been utilized the lymph content ratio between unaffected and affected lymphedema arms as a good meter to establish severity degree of the cancer-related lymphedema [4, 5]. Nevertheless, often these differences in electrical resistance between the affected and unaffected arms not reaches 10 Ohm, or a very few difference in considering that the volume increase in the affected limb could reach about 200 ml [5]. Another type of patients for which there may be differences in the water content between the two arms are those periodically submitted to hemodialysis [6]. In fact, by utilizing the multifrequency bioimpedance method, it has been found that, both prior and after dialysis, patients with right-side brachial fistulae when compared with the left side arm had a greater percentage of brachial extra cellular water (or lower electrical impedance at an injected current with a low frequency) with respect to the total body one. On the basis of what is described above, it appears that some measurement errors can occur if, before the clinical treatment, the impedance of the two arms turns out to be different. In fact, one might expect that in the dominant arm, due to its greater use, the muscular masses may be more consistent than in the auxiliary arm, and this would result in the presence of a greater quantity of water in that dominant arm, hence a lower value of the resistive component of its bioimpedance may occur. To verify if the above suggestion possesses or not a concrete foundation, in a group of healthy subjects we compared electrical resistance changes in both right and left upper arms to which was applied an alternated electrical current by means of a low constant voltage (1.0 Vpp) with a sweep of frequency from 1kHz to 300 kHz.

## II. Materials and methods

### A. Subjects

Twenty healthy subjects of which 9 males and 11 females, (age: 28.4 ± 1.4 years, weight: 63.8 ± 11.8 kg, height: 167.4 ± 7.5 cm) underwent to the experiment. All the participants were right-handed and were medical students who attended the internship at the medical department or postgraduate specialists in internal medicine, which were employed at the University Polyclinic of Cagliari. All of them were very skilled about the experimental tests conduction, nonetheless they signed their informed consensus and, in any case, the experiment was done in respect with what was stated in the Helsinki declaration of 2000.

### B. Instrumentation and experimental protocol

Upper arms electrical impedance spectroscopy (EIS) was assessed by means of an homemade portable device which implemented the electronic board AD5933 from Analog Devices Inc., USA [7] by using the electrodes bipolar configuration; it was powered by a USB gate of a laptop which, in turn, was not connected to the fixed electricity network. The device injected an electrical current with a frequency sweep from 1 kHz to 300 kHz at a constant tension of 1.0 Vpp, so the maximum output current is 0.25 mA (typical Short-Circuit Current to Ground at VOUT) [7]. As shows figure 1, disposable ECG electrodes were used to inject/detect the EIS signals in the forearms.

**Fig. 1.**
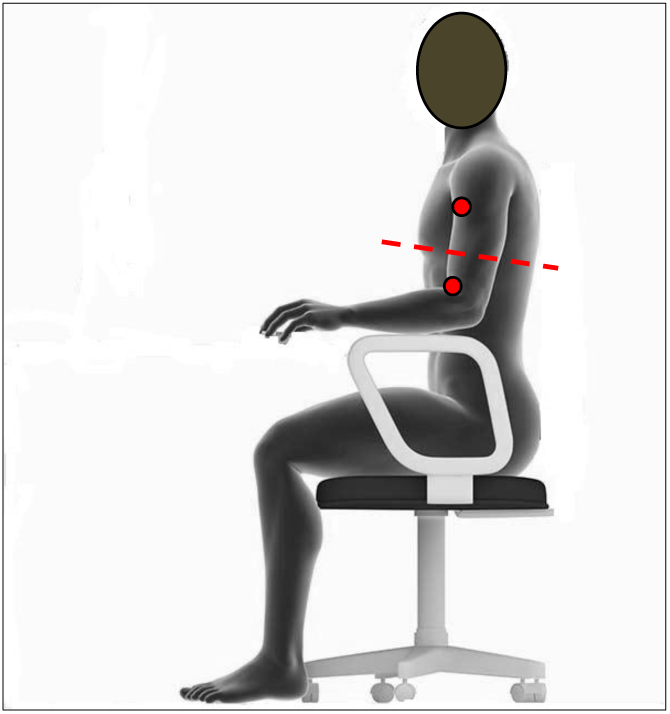
A male seated dummy of typical tested subjects is shown. Red circles represent bipolar connection of an upper arm to the EIS device and the dashed red line indicates the relative position of the biceps belly with respect to the atrioventricular septum.

The experimental sessions were made at the internal medicine ambulatory of the University Polyclinic of Cagliari, in Italy. Firstly, a medical history was acquired regarding possible pathological conditions that, being in place, could affect the quality of the bioimpedance information assessing. Then, age, weight and height of each subject were obtained as well as the length of each humerus bone that was made by an expert physician starting from the gleno-humeral joint until the olecranon fossa in the humerus bone. Also the maximal circumferences at the relaxed forearm muscles were measured just at the level of the biceps muscle belly.

The experimental sessions were made at the internal medicine ambulatory of the University Polyclinic of Cagliari, in Italy. Firstly, a medical history was acquired regarding possible pathological conditions that, being in place, could affect the quality of the bioimpedance information assessing. Then, age, weight and height of each subject were obtained as well as the length of each humerus bone that was made by an expert physician starting from the gleno-humeral joint until the olecranon fossa in the humerus bone. Also the maximal circumferences at the relaxed forearm muscles were measured just at the level of the biceps muscle belly.

After these preliminary anthropometric measurements were done, each subject was sitting on a comfortable chair with the upper body muscles as relaxed as possible and with each hand resting on the quadriceps muscle of the same side leg and having the palm facing upwards. The elbow angle of each arm was carefully chosen by the operator in such a way to approximate at the best the biceps belly position on the ideal line on which lies the atrioventricular septum of the heart. As is known [8], in standing subjects the spatial position of this heart septum, referred to the ground, corresponds to the condition in which the hydrostatic component of blood pressure is practically equal to zero. In this way, by placing the two electrodes equidistantly from the biceps belly along the humerus, the transmural pressure of the capillary vessel wall, which affects the flow of water and solutes between the inner and outer of blood capillaries, practically depended solely on the vis a tergo generated by the systolic ejection of the heart, thus minimizing extra-vascual water accumulation or its reduction due to adding or subtracting the hydrostatic component to the blood pressure generated by the cardiac systole. Obviously, the reaching of this hemodynamic equilibrium through the measurement points of the arms will consent to better assess possible differences in the water content due to physiological or pathological differences between forearms that took place previously, thanks to the comparison among the real components, i.e. the electrical resistance (R_F_), of the arms impedance. Then, in each forearm two disposable electrodes were applied just in correspondence of upper and lower tendon-muscle junctions making sure that both were equidistant from the biceps belly (see Figure 1). After a few minutes in which upper body of the subject was as relaxed as possible so that the only muscular force present in the muscles of the forearms was that depending from their basic postural control, a condition this latter that was semiotically verified by a skilled physician through specific palpations of the arms, then the EIS measurements could begin.

Each EIS measurement was repeated 3 times and the mean lasting time measurement for an electrode couple was about 20 s.

### C. Instrumentation and experimental protocol

About the impedance variables assessed by the board AD5933 from each forearm, we considered only the real component of the arms impedance, i.e. the R_F_, since its magnitude well represents the water volume within a biological tissue. From each arm of each subject we acquired R_F_ values corresponding to increasing frequency steps of 15 kHz (f_step_), starting from 15 kHz. We chose this relatively high frequency value in order to avoid noise and distortions in the acquisition of the electrical signals that are typical of low frequency circuits [9]. Signals from arm electrodes were acquired by the AD5933-development and this data was exported to laptop PC with a dedicate software. The software processed the acquired data in such a way of obtain the R_F_ mean value of those three assessments in succession by the device from each forearm of each subject at each increasing fstep. Then the cumulative mean ± SD was calculated among the previous calculated mean R_F_ values of each subject at a given fstep, making sure to separate data from right forearms to those from the left ones. To verify if possible differences among compared data were statistically significant the Student’s t test for paired data was applied, and a P < 0.05 was considered as significant. Where appropriate, we applied the regression method to assay possible mathematical function which could tie some variables together. Moreover, the Bland and Altman plot [10] was also applied to investigate on relationships among possible right versus left forearms differences in R_F_ occurring while fstep increased. In fact, this method compares two different generator of the same variable which, in this case, where the right and left forearms that both gave the same variable: the R_F_. The hypothesis was that if the two forearms were structurally equals then the assessed R_F_ from both of them could be identical at each applied frequency. Otherwise the structure of the arms would have been different and this, above all, would be due to the different quantity of water into them. The MedCalc statistical package (Belgium) was utilized to investigate about data differences.

## III. Results

Anthropometric measurements of forearms showed that no statistically significant difference existed between right (28.6 ± 2.6 cm) and left (28.7 ± 2.7 cm) homerus length (P = 0.81) as well as between right (27.5 ± 3.5 cm) and left (27.6 ± 3.3 cm) circumference of the forearms at level of the biceps belly (P = 0.59). Thanks to those anthropometric results which, reasonably, allowed us to consider that between the forearms of the subjects tested there were no morphological differences, we avoided to normalize assessed values of resistance with the forearms dimensional parameters. As shown in Table 1, we compared differences in R_F_ between forearms only in correspondence of discrete intervals of the frequency sweeping, i.e. the fstep.

**Table 1.** Mean vales ± SD among tested subjects are represented concerning resistance values respectively of right and left forearm assessed at each established step of current frequency. *R*_*F r-l*_ are the differences in the forearms mean resistance at each frequency step. (*) P < 0.05 for the marked *R*_*F r-l*_.

| Frequency [Hz] | Forearms Resistance [Ohm] |  |  |
| --- | --- | --- | --- |
| | Right | Left | $R_{F-r,l}$ |
| 15000 | $631.7 \pm 88.3$ | $659.2 \pm 88.5$ | -27.5* |
| 30000 | $550.3 \pm 65.5$ | $569.4 \pm 85.4$ | -19.1* |
| 45000 | $525.8 \pm 73.7$ | $535.9 \pm 80.4$ | -10.1 |
| 60000 | $518.3 \pm 62.4$ | $524.9 \pm 72.3$ | -6.6 |
| 75000 | $516.3 \pm 53.8$ | $521.4 \pm 64.7$ | -5.1 |

The table 1 indicates that, with the fstep that increased from 15 kHz to 45 kHz, both the right and left values of R_F_ reduced progressively. Interestingly, the mean value of right forearm R_F_ was significantly lower than in the left one.

Nevertheless, as the frequency increased the R_Fr-l_ reduced progressively up to became not statistically significant when the injected current overcame a frequency of 45 KHz. So we continued to assess R_F_ values only up to a fstep of 75 kHz. Figure 2 shows the regression curve of the R_Fr-l_ values versus the corresponding f_step_ and it is well represented (P = 0.003) by the equation (1).

**Fig. 2.**
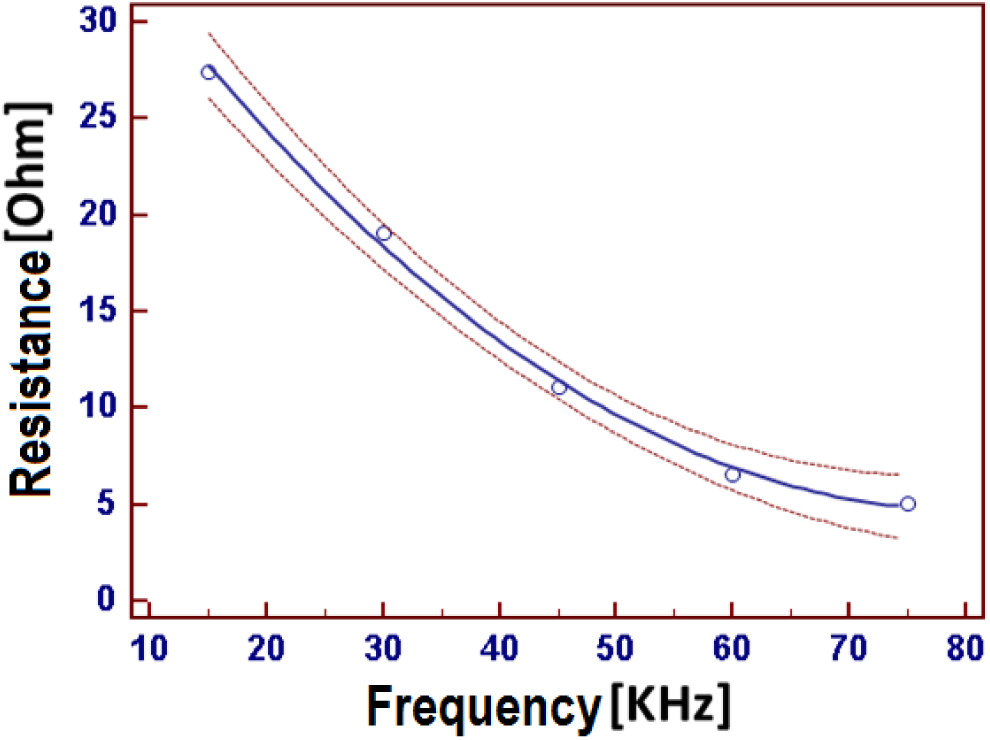
Blue line represents the branch of a descending parabola concerning the quadratic regression of the differences between right and left resistance values (Ohm) versus the corresponding frequency step (KHz) of the applied current to each forearm. Empty circles are the real values, red dashed lines represent 95% of the confidence interval.

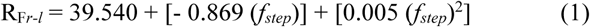

Equation (1) consents to quantify the behaviour of the R_Fr-l_ when the fstep increased. In fact, it appears that if on the one hand the R_Fr-l_ would increase almost in a unitary way when the fstep increases by each expected step of 15 kHz (as shown by the first term of the equation in which fstep has the exponent 1), at the same time this increase of R_Fr-l_ is powerful contrasted by the second term of the equation where fstep has exponent 2. In fact, stepping from 15 to 30 kHz the reduction in R_Fr-l_ was of 8.3 Ω while stepping from 60 to 75 kHz the R_Fr-l_ reduced of only 1.5 Ω.

The Bland & Altman plots represented in figure 3 show that, from 15 kHz to 60 kHz, the R_F *r-l*_ of quasi all the tested subjects, plotted against the averages of the two measurements, fall within the 95% confidence interval. The plots also shows that the gap between the X axis corresponding to zero differences (red lines) and the parallel line to the X axis that corresponds to the real mean of the R_F*r-l*_ (black lines) are the mean bias between the two forearms. Also in the plots is shown that, after the frequency step of 30 kHz, the regression line of the R_F*r-l*_ values versus the measurements averages is progressively inclined downwards to right.

**Fig. 3.**
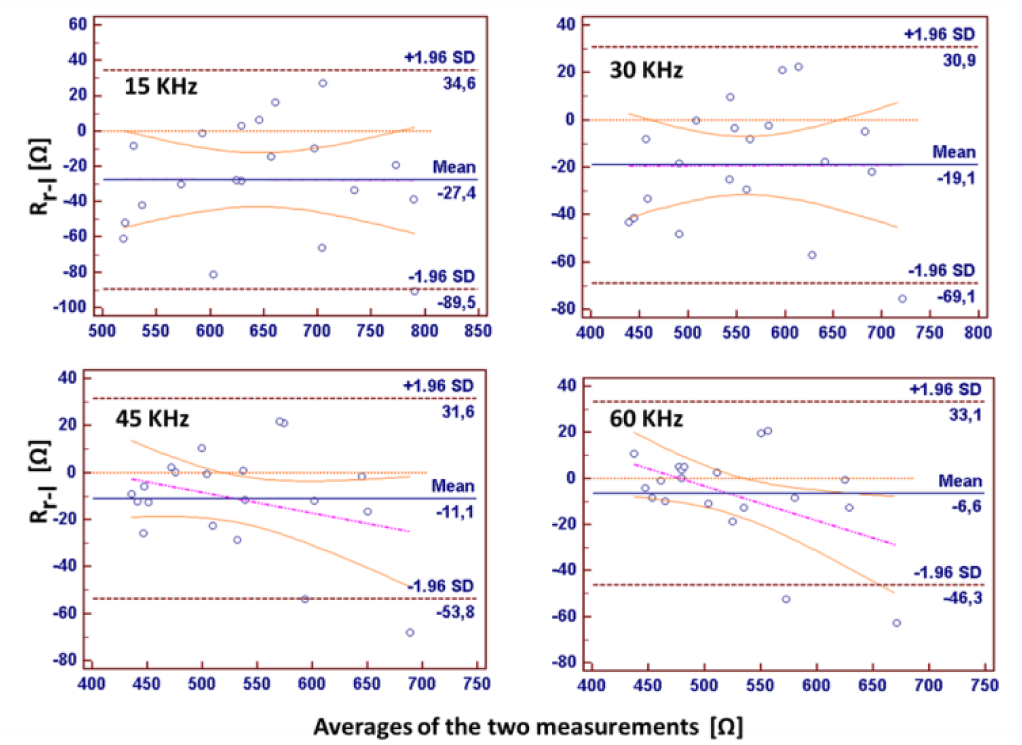
Bland and Altman plots are presented for frequency step increasing from 15 to 60KHz. Each empty circle represent a tested subject. They show that the gap between the X axis, corresponding to zero differences (red straight line) and the parallel line to the X axis that concerns to the real mean of the RFr-l (black straight line) didn’t overlap. Moreover, the line of the regression among points shows to reduce its pendency when frequency overcame 45 KHz.

## IV. Discussion

In summarizing the results from these experiments we could highlight three main occurrences. The first concerns the lower R_F_ values assessed from the right forearm with respect to the left one; the second refers to the progressive reduction in this R_F_ right versus left difference as the current frequency increases; the third is the tendency to reduce more the right R_F_ with respect the left one in those subjects who showed highest average values in the two measurements when the frequency was over 30 kHz.

The first occurrence could be explained considering that the main arm could have developed greater muscle mass at the level of the biceps due to the greater frequency and strength with which this muscle is daily engaged compared to that of the left arm. The increase in muscle mass of the biceps agrees well with a consequent increase in the intra and extracellular water content also due to the fact that the prolonged physical training of the main arm could induce a significant increase in its vascularisation [11]. Therefore, larger volume of water into the right forearm well justify its lover R_F_ with respect the left one. Concerning the second occurrence for which R_F_ progressively diminished with the frequency increase, it could be considered that, at relatively low frequencies, the current could be blocked by membrane capacitances that could stay serially positioned with respect to resistive components through which it was flowing. As the frequency increased those capacitances behaved as short-circuit thus unlocking those resistive component serially connected to them, so enlarging the number of resistive components staying together in parallel [12]. Then in both forearms, the observed frequency dependent, progressive reduction of the R_F_, could be justified in this way. As shows the Bland and Altman diagrams, the R_F*r-l*_ increased progressively at highest frequencies as increased the averages of the two measurements. This could be explained admitting that, in the dominant arm, the intracellular organelles were more and bigger than in the auxiliary one, due to the higher physical engagement of this arm. These organelles could be considered as cells inside the cells, i.e. an electrolytic medium contained in a dielectric membrane immerse into an electrolytic medium, for this, the membranes of organelles were in turn short-circuited by very high frequency so unlocking the current passage through they [12]. This effect, in the cells of the main arm could result in an increasing fall of the R_F_, as a function of the increasing frequencies, which was more consistent than in the auxiliary arm. As the consequence of that occurrence, a progressive R_F*r-l*_ reduction depending from the augmenting frequency of the injected current might happens.

## V. Conclusions

These experimental results highlights the risk of the occurrence of some mistakes when lymphedema may occur in one arm and EIS was utilized to monitoring, in that arm, the water volume trend in comparison with the other side arm. In fact, these results underline in the main forearm a largely low value of the resistance than in the auxiliary one, even in healthy subjects. So, care must be taken when the EIS was adopted in these clinical assessments. A further suggestion arising from these results concerns the identification of a low threshold in the injected current frequency since they shows that down 45 kHz the resistance differences among main and auxiliary arm could be present.

## Acknowledgment

*The Authors thanks the Italian Ministry of Foreign Affairs and International Cooperation (MAECI), which granted a scholarship to an Argentinian researcher who carries out research in co-tutoring in Cagliari -Italy, thanks to which it has been possible to obtain these experimental results*.

## References

1. I. Rogowski, T. Creveaux, C. Genevois, S. Klouche, M. Rahme, P. Hardy, Upper limb joint muscle/tendon injury and anthropometric adaptations in French competitive tennis players. Eur. J. Sport Sci., vol. 16, pp. 483–489, 2016.

2. R. J. Tsai, L. K. Dennis, C. F. Lynch, L. G. Snetselaar, G. H. D. Zamba, C. Scott-Conner, Lymphedema following breast cancer: The importance of surgical methods and obesity. Front. Womens Health, Jun;3(2), 2018.

3. D. M. Kaulesar Suku, P. T. den Hoed, E. J. Johannes, R. van Dolder, E. Benda, Direct and indirect methods for the quantification of leg volume: comparison between water displacement volumetry, the disk model method and the frustum sign model method, using the correlation coefficient and the limits of agreement. J. Biomed. Eng., vol. 15, pp 477–480, 1993.

4. A. G. Warren, B. A. Janz, S. A. Slavin, L. J. Borud, The use of bioimpedance analysis to evaluate lymphedema. Ann. Plast. Surg., vol. 58, pp. 541–5433, 2007.

5. B. Smoot, S. Zerzan, J. Krasnoff, J. Wong, M. Cho, M. Dodd, Upper extremity bioimpedance before and after treadmill testing in women post breast cancer treatment. Breast Cancer Res. Treat., vol. 148, pp. 445–453, 2014.

6. S. Kumar, M. Khosravi, A. Massar, M. Potluri, A. Davenport, Changes in upper limb extracellular water content during hemodialysis measured by multi-frequency bioimpedance. Int. J. Artif. Organs, vol. 36, pp. 203–207, 2003.

7. F. Noveletto, P. Bertemes-Filho, D. Dutra, Analog Front-End for the Integrated Circuit AD5933 used in Electrical Bioimpedance Measurements, in: Book of 2° Latin American Conference on Bioimpedance, pp. 48–51, 2016.

8. R. B. Berne, M. N. Levy, Cardiovascular Physiology, McGraw-Hill, New York, 1988.

9. D. Bucci, Introduction to noise analysis in low frequency circuits, in: Analog electronics for measuring systems, Ed. D. Bucci, ISTE ltd, London, pp. 121–152, 2017.

10. D. Giavarina, Undertanding Bland Altman analysis, Biochemia Medica, vol. 25, pp. 141–151, 2015.

11. T. L. Haas, E. Nwadozi, Regulation of skeletal muscle capillary growth in exercise and disease, Appl. Physiol. Nutr. Metab., vol. 40, pp. 1221–1232, 2015.

12. A. Ivorra, Bioimpedance monitoring for physicians: an overview, https://www.researchgate.net/publication/253563215_Bioimped-ance_Monitoring_for_physicians_an_overview.

